# Two canonically aerobic foraminifera express distinct peroxisomal and mitochondrial metabolisms

**DOI:** 10.1101/2022.07.20.500910

**Authors:** Christopher Powers, Fatma Gomaa, Elizabeth B. Billings, Daniel R. Utter, David J. Beaudoin, Virginia P. Edgcomb, Colleen M. Hansel, Scott D. Wankel, Helena L. Filipsson, Ying Zhang, Joan M. Bernhard

**Author notes:** Correspondence: Ying Zhang, Joan M. Bernhard.

## Abstract

Certain benthic foraminifera are known to thrive in marine sediments with low oxygen or even without detectable oxygen. Potential survival avenues used by these supposedly aerobic protists include fermentation and anaerobic respiration, although details on their adaptive mechanisms remain somewhat elusive. To better understand the metabolic versatility of foraminifera, we studied two benthic species that thrive in oxygen-depleted marine sediments. Here we detail, via transcriptomics and metatranscriptomics, differential gene expression of *Nonionella stella* and *Bolivina argentea*, collected from Santa Barbara Basin, California, USA, in response to varied oxygenation and chemical amendments. Organelle-specific metabolic reconstructions revealed that these two species utilize adaptable mitochondrial and peroxisomal metabolism that reflect their differing lifestyles. *N. stella*, most abundant in anoxia and characterized by the lack of food vacuoles and the abundance of intracellular lipid droplets, was predicted to couple the putative peroxisomal beta-oxidation and glyoxylate cycle with a versatile electron transport system and a partial TCA cycle running in the reductive direction. In contrast, *B. argentea*, most abundant in hypoxia and contains food vacuoles, was predicted to utilize the putative peroxisomal gluconeogenesis and a full TCA cycle but lacks the expression of key beta-oxidation and glyoxylate cycle genes. These metabolic adaptations likely confer ecological success while encountering deoxygenation and illuminate the importance of metabolic modifications and interactions between mitochondria and peroxisomes in protists.

**Importance:** Foraminiferan protists are nearly ubiquitous in today’s oceans and likely were major components of the Neoproterozoic protistan community. While largely considered aerobic, certain foraminifera demonstrate surprising adaptability to hypoxia and anoxia, contributing to biogeochemical cycling in benthic environments. The analyses of Rhizarian adaptive metabolism set the stage for studying other microeukaryotes under increasing ocean deoxygenation. Revealing the metabolic roles of foraminifera in anaerobic biogeochemical cycling should spur reassessments of existing paleoecological datasets as well as new perspectives on the metabolic evolution of eukaryotic cells.

## Introduction

Foraminifera are widely distributed across the marine realm, where they can constitute large proportions of the benthos (1, 2). This diverse protistan group is broadly implicated in biogeochemical cycling, where some foraminifera and/or their symbionts perform approximately 67-100% of total denitrification (3–5). Others are involved in sulfur cycling (6) and reactive oxygen species (ROS) chemistry (7). Certain foraminiferan species, while exhibiting lower abundance in oxygenated habitats, thrive in hypoxic or anoxic sediments that are characterized by low oxygen or absence of dissolved oxygen, respectively (8–10).

Robust foraminiferan populations occur across the oxygen gradient within chemoclines. Chemoclines are often coupled with complex gradients of reduction-oxidation chemistry (11) and considerable concentrations of hydrogen sulfide (8). Reactive oxygen species, such as oxygen radicals and hydrogen peroxide, can be produced in chemoclines via abiotic means such as the interaction of O_2_ with reduced metals, sulfide, or dissolved organic carbon, or through biotic means mediated by transmembrane NADPH oxidases (12). Certain chemocline-associated foraminifera contain masses of peroxisomes, often correlated with the presence of endoplasmic reticulum (7). It has been hypothesized that peroxisomes can mediate foraminiferal aerobic respiration through catalase-induced O_2_ production (7), but this hypothesis requires further investigation.

Protists inhabiting hypoxic or anoxic environments have been shown to modify the functions of their mitochondria and peroxisomes (13–18). The mitochondria related organelles (MROs), a reduced form of mitochondria characterized by the loss of key enzymes involved in oxygen-dependent metabolism, are often specialized for anaerobic energy generation through pyruvate fermentation, hydrogen production, substrate-level phosphorylation, and malate dismutation (13, 14, 19), or known to abandon their roles in energy metabolism entirely (15, 19, 20). The metabolism of MROs can be evolutionarily linked to metabolic modifications of peroxisomes. In some obligate anaerobic protists, the peroxisomes are often absent, as observed in *Diplonema papillatum, Entamoeba histolytica, Pelomyxa schiedti,* and *Mastigameoba balamuthi* (17, 18, 21, 22). Other obligate anaerobes may retain peroxisomes, but lose the O_2_-involving catalase and beta-oxidation functions (17, 18, 22).

The known metabolic capacity in benthic foraminifera may include denitrification (4, 23, 24), ammonium assimilation (23, 25), fermentation (23, 26), and utilization of dissolved organic matter (26, 27). Our recent findings, based on transcriptomic and metatranscriptomic analysis of two foraminiferan species, *Nonionella stella* and *Bolivina argentea*, collected from the oxygen-depleted Santa Barbara Basin (SBB), USA, indicate that both employ anaerobic pyruvate metabolism and have the ability to perform denitrification for survival in anoxia (23). Despite these shared metabolic traits, the two species inhabit distinct niches within the SBB (Table 1). *N. stell*a is most abundant in severely hypoxic to euxinic (anoxic plus sulfidic) conditions, and is the sole calcareous foraminifer documented to survive long-term anoxia within the basin (9, 28). *N. stella* sequesters chloroplasts (23, 29, 30), likely from prey, proliferates peroxisome-endoplasmic reticulum (P-ER) complexes (7), and apparently lacks food vacuoles (23). In contrast, *B. argentea* is abundant at the basin’s flanks, rarely occurring in the most oxygen-depleted conditions (9), contains solitary to small groups of peroxisomes, and exhibits evidence of digestion of chloroplasts and other organics (5). Such contrasting physiological features make these two species ideal for examining the metabolism of benthic foraminifera that are well adapted to the variable biogeochemistry known to occur episodically within the SBB (8, 11, 31). The metabolic function of intracellular organelles has not been elucidated in the two foraminifera, leaving potential knowledge gaps in the organelle responses to environmental variability. Here we apply metabolic reconstructions of both species to reveal potential metabolic connectivity of modified peroxisome and mitochondrion that allow these foraminifera to thrive in differing concentrations of oxygen and varied redox chemistry. Our study suggests distinct metabolic responses by *N. stell*a and *B. argentea* to the addition of hydrogen peroxide (H_2_O_2_), nitrate (NO_3_^-^), and/or hydrogen sulfide (H_2_S) amendments under anoxia or hypoxia, providing insights into the metabolic adaptations of foraminifera to chemocline biogeochemistry.

## Results

### Production and maintenance of peroxisomes

Genes related to the function and biogenesis of peroxisomes were identified in the (meta)transcriptomes of *N. stella* and *B. argentea* (Figure 1). Specifically, transcripts were found encoding a transmembrane protein (Pex11) linked to remodeling of the peroxisome membrane for proliferation of peroxisomes (32, 33), transport of fatty acids for peroxisomal beta-oxidation (34), and initiation of contact between mitochondria and peroxisomes (35). Transcripts of three receptor proteins (i.e. Pex5, Pex7, Pex19) that recognize signals for peroxisomal protein import were also identified (36). Further, expression of genes encoding AAA ATPases (i.e. Pex1, Pex6) and ubiquitination proteins (i.e. Pex4, Pex2, Pex10) demonstrated activities in the release of peroxisomal imported proteins from signal receptors and the recycling of receptor proteins, respectively (36). It is notable that transcription of all the *Pex* genes was observed in *N. stella*, while transcription of the *Pex2*, *Pex11*, and *Pex19* genes was not identified in *B. argentea* under any treatment (Figure 1 **& SI Dataset S1**).

**Figure 1:**
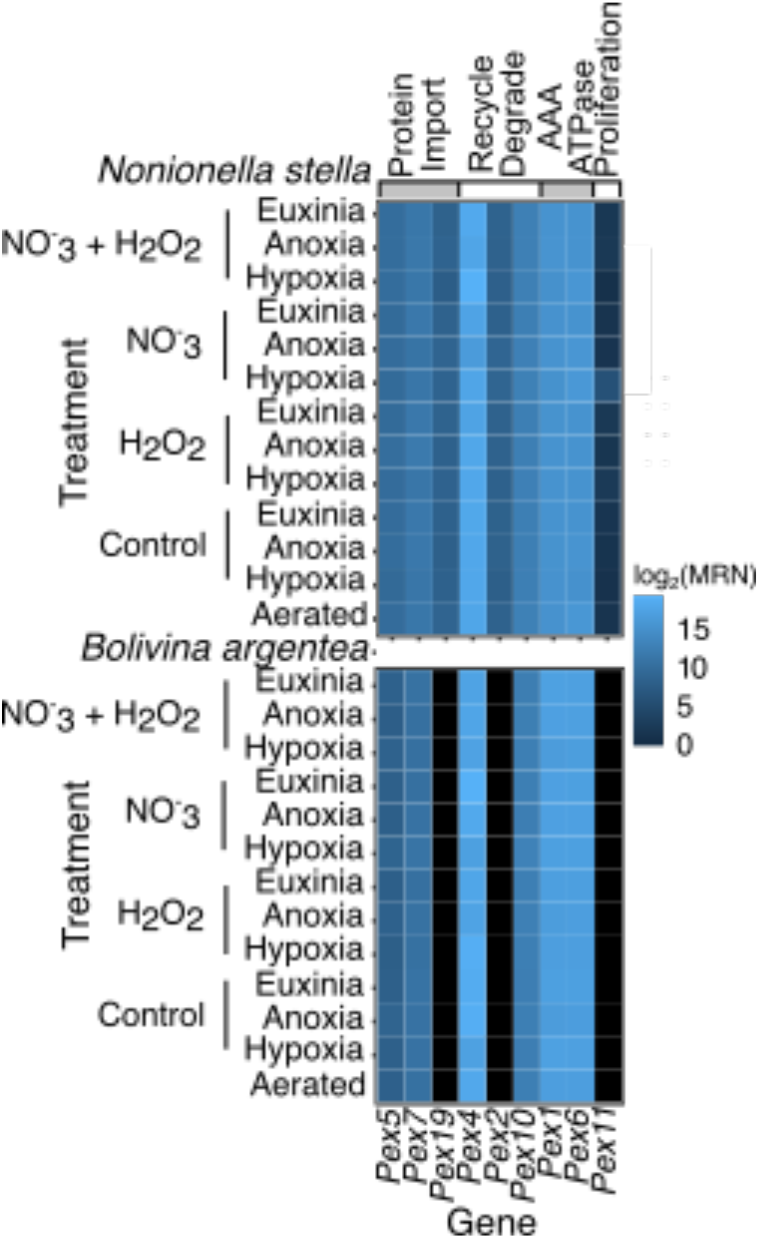
Heatmap presented by species showing the log_2_ transformed median ratio normalization (MRN) values for all the identified peroxisome biogenesis genes. Black indicates no expression.

### Metabolism of the peroxisomes and mitochondria

A reconstruction of metabolic pathways, based on transcripts identified in the (meta)transcriptomes of *N. stella* and *B. argentea,* revealed an overall distinction in the metabolism of the two species (Figure 2). Protein localizations were predicted based on the consensus of multiple approaches for the characterization of peroxisomal and mitochondrial functions (**Materials and Methods**), and the results are summarized below.

**Figure 2:**
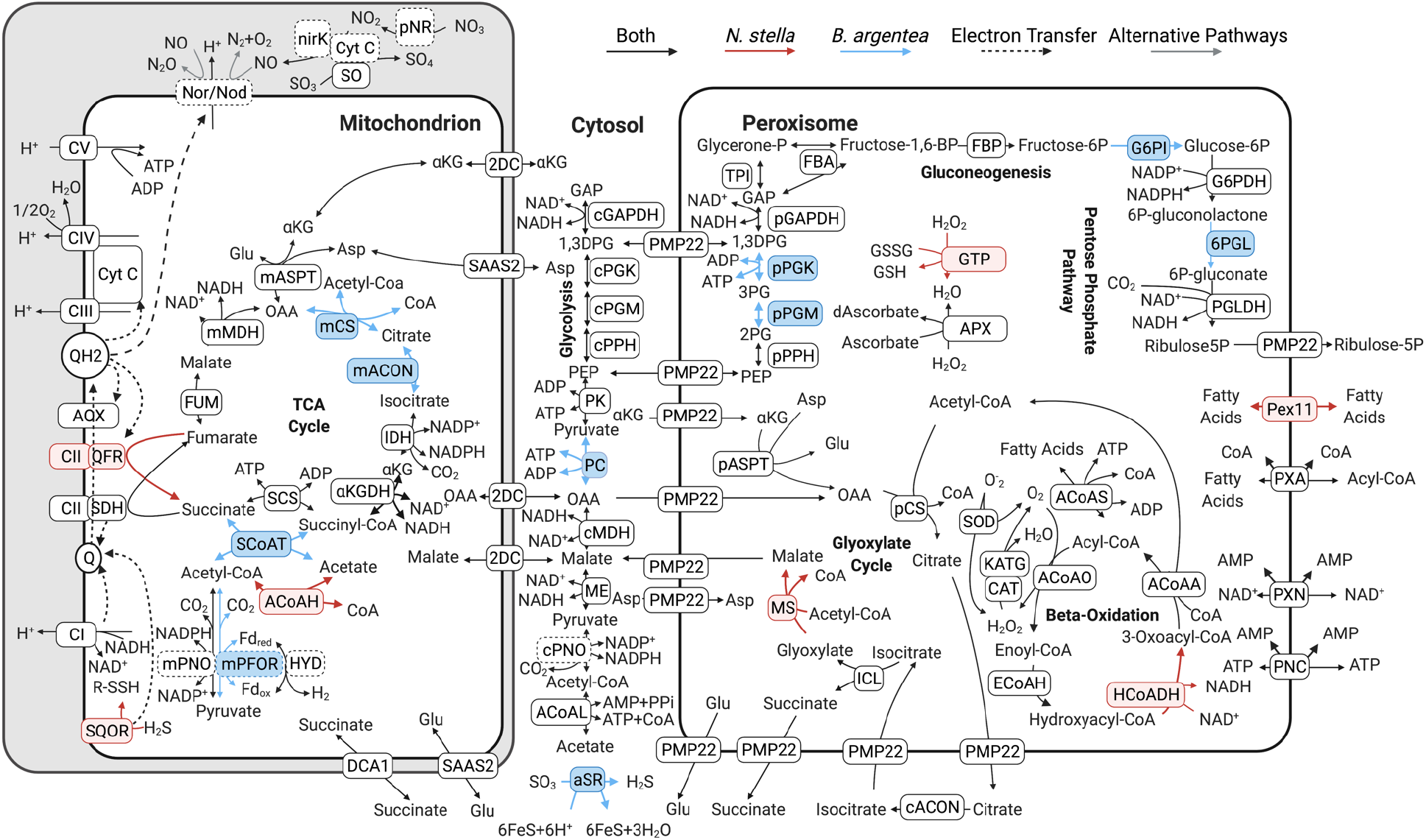
Pathway map of the peroxisomal and mitochondrial metabolic processes predicted in *N. stella* and *B. argentea*. Unique genes expressed in *N. stella* are shown with pink shade with the corresponding metabolic reactions indicated as red arrows. Unique genes expressed in *B. argentea* are shown with blue shade with the corresponding metabolic reactions indicated as blue arrows. Genes expressed in both species are shown in black boxes with the corresponding metabolic reactions indicated as black arrows. Biochemical conversions between metabolites are annotated with the abbreviation of corresponding genes, as defined in **SI Dataset S1**. Boxes with a dotted outline indicate genes that have been described previously (23). Gray arrows represent alternative pathways for the same genes, specifically with respect to the currently uncertain role of Nor/Nod (23). Q represents quinones/rhodoquinones and QH2 represents quinols/rhodoquinols. Cyt C represents Cytochrome C. CI, CII, CIII, CIV, and CV, represent Complex I, II, III, IV, and V, respectively, of the electron transport chain. Dashed arrows indicate the flow of electrons.

Proteins involved in beta-oxidation and the glyoxylate cycle were predicted to localize in the peroxisome. Beta-oxidation, which has not been previously described in foraminifera, is linked to the detoxification of H_2_O_2_ or syperoxide radicals by catalase/peroxidase (e.g. CAT, KATG) or superoxide dismutase (SOD), respectively. Beta-oxidation breaks down fatty acids for the production of acetyl-CoA, which in turn serves as a precursor metabolite for the glyoxylate cycle, leading to the production of central carbon metabolites, such as malate and oxaloacetate (OAA) (Figure 2). All genes associated with the above-mentioned peroxisomal pathways were expressed in *N. stella*, while transcripts that encode 3-hydroxyacyl-CoA dehydrogenase (HCoADH) of the beta-oxidation pathway and malate synthase (MS) of the glyoxylate cycle were not identified in *B. argentea* across all conditions examined (Figure 3). Similarly, genes encoding Pex11, which supports the transport of fatty acids for enabling beta-oxidation in the peroxisomes (34), were expressed in *N. stella* but not in *B. argentea* (Figure 1).

**Figure 3:**
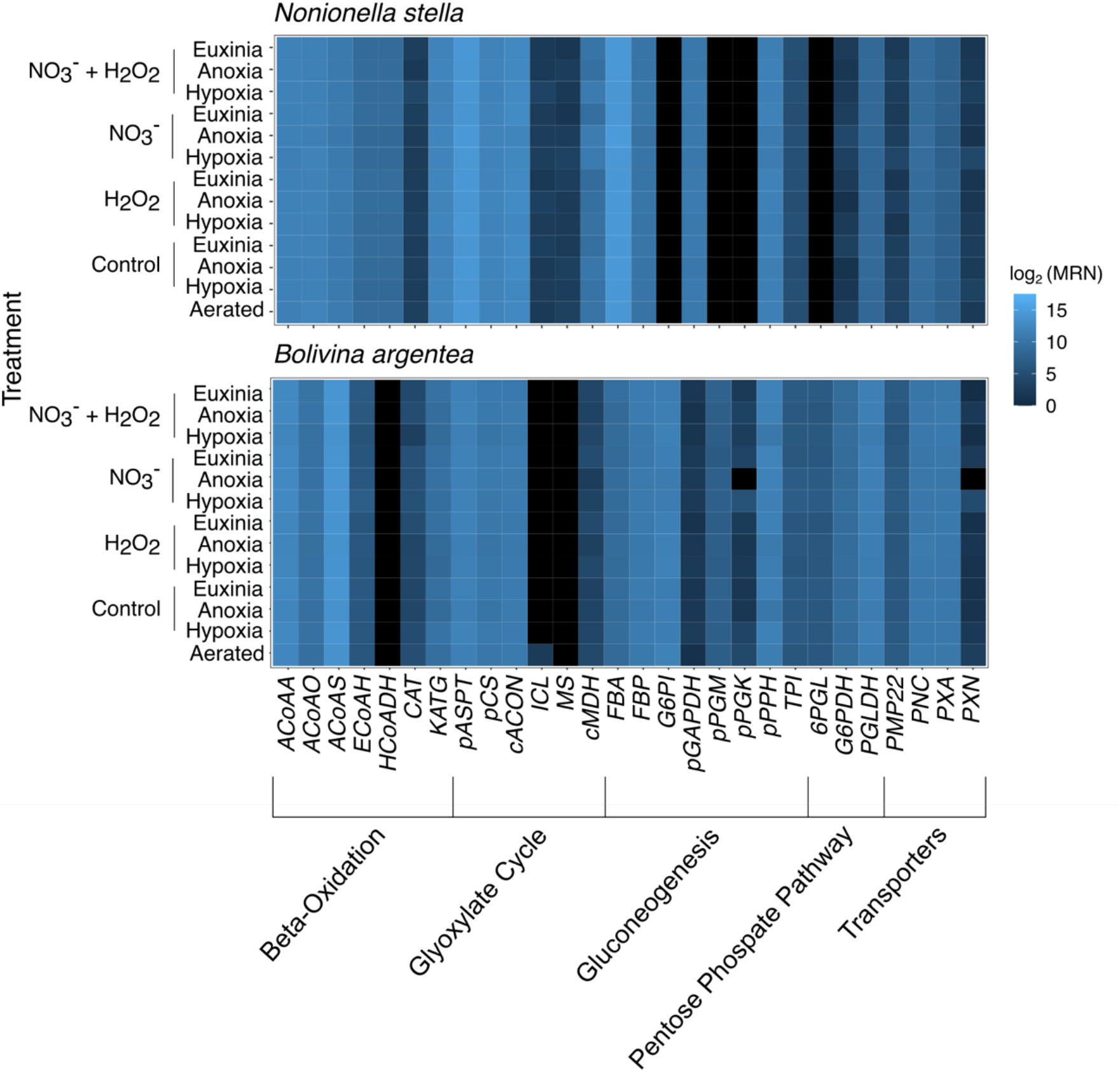
Heatmap showing the log_2_ transformed median ratio normalization (MRN) values for all genes that encode peroxisomal proteins of each species. Black indicates no expression. Abbreviations align with those in **SI Dataset S1**.

Transcripts encoding fructose bisphosphatase (FBP), a committed step of gluconeogenesis, were identified and predicted to localize in the peroxisome of both *N. stella* and *B. argentea*. However, the predicted peroxisomal gluconeogenic pathways in the two species had distinct entry points using metabolites from the cytosol: 1,3-bisphospho-D-glycerate (1,3DPG) for *N. stella*; phosphoenolpyruvate (PEP) for *B. argentea* (Figure 2). Similarly, the oxidative pentose phosphate pathway were predicted to localize in the peroxisome of both species, but it appeared to be active in *B. argentea* but not in *N. stella* because *N. stella* lacked expression of genes encoding the peroxisome-localized 6-phosphogluconolactonase (6PGL) or glucose-6-phosphate isomerase (G6PI) (Figures 2 **&** 3).

The prediction of protein localization in mitochondria also revealed distinct transcription of metabolic pathways in *N. stella* and *B. argentea*. Expression of genes for mitochondrial substrate-level phosphorylation was observed in both species, with transcripts encoding acetyl-CoA hydrolase (ACoAH) identified in *N. stella* but not *B. argentea*, and transcripts encoding succinate:acetate CoA transferase (SCoAT) found in *B. argentea* but not *N. stella.* Related to the utilization of acetyl-CoA, genes encoding pyruvate:NADP oxidoreductase (mPNO) were expressed in both species and genes encoding pyruvate:ferredoxin oxidoreductase (mPFOR) were expressed in *B. argentea*. Further, transcripts encoding the sulfide-oxidizing sulfide:quinone oxidoreductase (SQOR) were detected in *N. stella*, but not in *B. argentea* (SI Figure S1). Finally, a complete mitochondrial tricarboxylic acid (TCA) cycle was identified from the (meta)transcriptomes of *B. argentea*, while only a truncated TCA cycle, from oxaloacetate (OAA) to succinyl-CoA, was detected in the *N. stella* (meta)transcriptomes (Figure 2).

Core subunits of the electron transport chain (ETC) Complexes I-V were identified in both species. However, variations were observed in the subunit compositions of Complex II (CII) (SI Figure S1). Specifically, characterization of succinate dehydrogenase (SDH) was based on the overall sequence similarity to experimentally verified reference proteins and the presence of an isoleucine and other common variants in the quinone-binding site of the large cytochrome subunit (CII-SDHC) (37). The characterization of quinol:fumarate oxidoreductase (QFR) was based on the presence of a glycine in a key position of the quinone-binding pocket, which indicated an essential modification in the QFR large cytochrome subunit (CII-QFRC) for the binding of rhodoquinone, an electron carrier required for anaerobic energy metabolism through fumarate reduction (37, 38) (Figure 4A). While transcripts encoding the CII-SDHC were detected in both *N. stella* and *B. argentea*, putative transcripts of the CII-QFRC were identified solely in *N. stella* (Figure 4B). The putative CII-SDH and CII-QFR formed a monophyletic group with other foraminifera (e.g. *Rosalina*, *Reticulomyxa*, *Sorites*, *Elphidium*, *Ammonia*). Presence of the same QFR-conserved glycine mutation was observed among other taxa known to exhibit anaerobic metabolism (i.e. *Mastigamoeba balamuthi*, *Blastocystis hominis*, and the metazoans *Ascaris suum*, *Caenorhabditis elegans*, *Ancylostoma ceylanicum*) (Figure 4A). The biosynthesis of rhodoquinone was putatively identified in both *N. stella* and *B. argentea* based on the detection of transcripts encoding rhodoquinone biosynthesis protein A (RQUA) (Figure 4B), which catalyzes the conversion of ubiquinone to rhodoquinone (39). Multiple sequence alignment revealed a conservation of quinone binding site residues between the RQUA proteins of foraminifera and other eukaryotes that perform fumarate reduction during exposure to anoxia (Figure 4C) (28).

**Figure 4:**
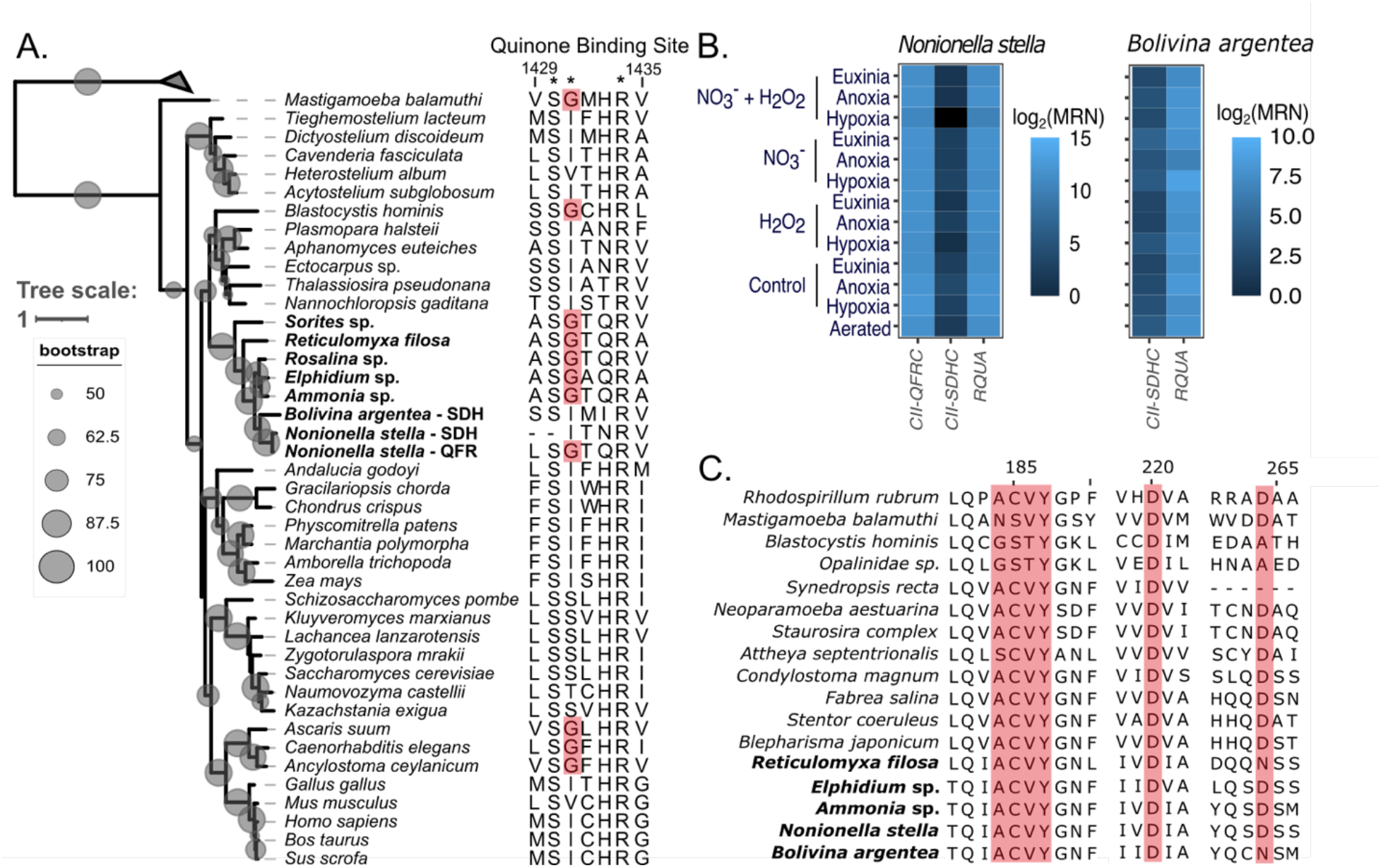
**A.** Phylogenetic reconstruction of concatenated protein sequences from Complex II showing phylogenetic placement of the putative CII-QFR and CII-SDH from *N. stella* and *B. argentea*, with bootstrapping support values represented as scaled, gray-shaded circles. This phylogeny is rooted using bacterial fumarate reductase sequences. Leaves belonging to foraminifera are highlighted with bold font. The amino acid sequences to the right of the tree shows an alignment of the quinone-binding site in the CII large cytochrome C subunit. Asterisks indicate amino acids implicated in quinone binding. Sites highlighted in red background are QFR-specific mutations associated with the binding of rhodoquinone. **B.** Expression of the *CII-QFRC*, *CII-SDHC*, and *RQUA* genes of *N. stella* and/or *B. argentea* represented as a heatmap for each species based on the log_2_ transformed median ratio normalization (MRN) values. **C.** Multiple sequence alignment of the RQUA protein from foraminifera and other eukaryotes. Amino acid residues highlighted in red background indicate functional sites in the quinone-binding pocket. Sequences belonging to foraminifera are labeled with bold font.

The mitochondrial electron transport system of both *N. stella* and *B. argentea* was complemented by alternative oxidase (AOX), which is known to mediate redox state of the quinone pool (40), and denitrification proteins (composed of pNR, NirK, and Nor/Nod) (23, 24). The *AOX* and denitrification genes were constitutively expressed in both foraminifers (**SI Dataset S1**)

Proteins that were not predicted to locate in any specific organelles were classified as cytosolic proteins. Transcripts encoding cytosolic proteins, such as malate dehydrogenase (cMDH), malic enzyme (ME), and acetyl-CoA ligase (ACoAL), were identified from the (meta)transcriptomes of both *N. stella* and *B. argentea* (Figure 2). Finally, transcripts encoding the cytosolic pyruvate carboxylase (PC) and assimilatory sulfite reductase (aSR) were identified in *B. argentea* but not in *N. stella* (Figure 2 **& SI Dataset S1)**.

### Differential gene expressions

*N. stella* responses to the addition of chemical amendments were revealed by the identification of differentially expressed metabolic genes under exposure to H_2_O_2_ and NO_3_^-^, particularly within the anoxic condition (**SI Dataset S2,** Figure 5). The H_2_O_2_ amendment produced no significant differential expression of genes compared to the anoxia control (without either H_2_O_2_ or NO_3_^-^ amendment) or the amendment of both H_2_O_2_ and NO_3_^-^. However, significant differential gene expression was observed between the H_2_O_2_ and the NO_3_^-^ only amendment (Figure 5A). Specifically, genes associated with denitrification (*pNR*, *NirK*, and *Nor/Nod*), sulfite oxidation (*SO*), and Complex V ATP synthase (*CV*) were up-regulated in the H_2_O_2_ amendment. In contrast, genes associated with subunits of Complex II (*CII*) were down-regulated in H_2_O_2_ compared to the NO_3_^-^ amendment. Genes related to several peroxisomal processes, such as the acyl-CoA importer (*PXA*), beta-oxidation of fatty acids (enoyl-CoA hydratase -*ECoAH* and 3-hydroxyacyl-CoA dehydrogenase - *HCoADH*), the glyoxylate cycle (*cACON* – cytosolic aconitase), gluconeogenesis (*FBP*), and *Pex7*, which encodes a receptor for peroxisomal matrix proteins, were also upregulated in the H_2_O_2_ amendment. Additional up-regulation of genes was observed among the cytosolic Enolase (*cPPH*) and glyceraldehyde-3-phosphate dehydrogenase (*cGAPDH*).

**Figure 5:**
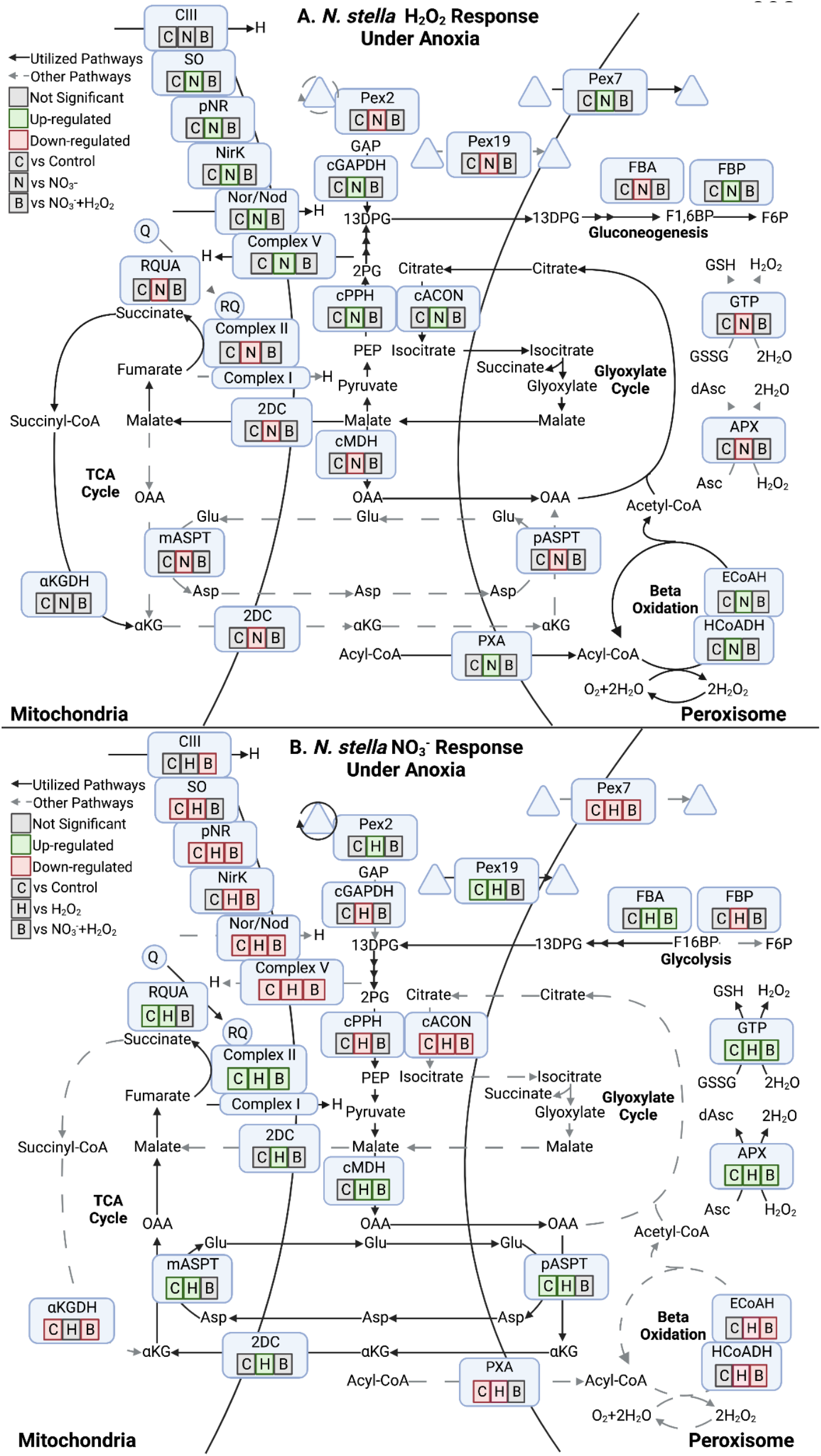
*N. stella* responses to chemical amendments under anoxia. **A.** Differential expression of genes under the H_2_O_2_ amendment in contrast to the control (C), NO_3_^-^ (N), and combined H_2_O_2_ + NO_3_^-^ (B) amendments. **B.** Differential expression of genes under the NO_3_^-^ amendment in contrast to the control (C), H_2_O_2_ (H), and combined H_2_O_2_ + NO_3_^-^ (B) amendments. Triangles in both panels represent transported and recycled proteins. Significant differential gene expressions between contrasts (adjusted p-value < 0.05) are highlighted in green (Up-regulated) or red (Down-regulated). Contrasting conditions with no significant differential gene expression are indicated in gray (Not Significant).

The transcriptional responses of *N. stella* to the addition of NO_3_^-^ under anoxia included the up-regulation of genes related to fumarate reduction (*CII* and *RQUA*) and the down-regulation of genes associated with denitrification (*pNR*, *NirK*, *Nor*/*Nod*) and Complex III (*CIII*) (Figure 5B). Several genes related to carbon metabolism, including the cytosolic malate dehydrogenase (*cMDH*), mitochondrial and peroxisomal aspartate transaminase (*mASPT* and *pASPT*), mitochondrial 2-oxodicarboxylate carrier (*2DC*) and peroxisomal glycolysis (fructose-bisphosphate aldolase - *FBA*), were also up-regulated in the NO_3_^-^ amendment. Lastly, the NO_3_^-^ amendment also stimulated up-regulation of genes associated with peroxisome biogenesis (*Pex19* and *Pex2*) and ROS response (ascorbate peroxidase - *APX* and glutathione peroxidase - *GTP*).

*N. stella* exhibited similar responses to the H_2_O_2_ and NO_3_^-^ amendments under hypoxia as compared to anoxia, but with modifications. The NO_3_^-^ amendment in hypoxia induced the up-regulation of *CII*, *CIV*, and genes encoding the peroxisome transmembrane protein Pex11 (SI Figure S3A). The combined amendment of both NO_3_^-^ and H_2_O_2_ induced the up-regulation of denitrification gene *NirK* and carbon metabolism genes, such as 2-oxoglutarate dehydrogenase (*aKGDH*), *cGAPDH*, and *cPPH* (SI Figure S3B).

Besides the H_2_O_2_ and NO_3_^-^ amendments, differential expression of metabolic genes was observed in *N. stella* in the presence or absence of oxygen and with or without the addition of H_2_S. For example, significant up-regulation of Complex IV (*CIV*) was identified in the aerated treatment compared to the hypoxic, anoxic, and euxinic treatments (SI Figure S2). Similarly, genes related to ROS responses, such as *APX*, *GTP*, and catalase-peroxidase (*KATG*), and the peroxisome biogenesis (*Pex19*), were up-regulated in the aerated condition (**SI Dataset S2**). Interestingly, the addition of H_2_S to the NO_3_^-^ amendment of anoxia (SI Figure S4) induced a similar response as the H_2_O_2_ amendment of anoxia (Figure 5A), with the up-regulation of genes for denitrification (*pNR*, *NirK*, *Nor*/*Nod*), beta-oxidation (*ECoAH* and *HCoADH*), the glyoxylate cycle (*cACON*), gluconeogenesis (*FBP*), and peroxisome receptor gene *Pex7* in the NO_3_^-^ amendment under euxinia.

In contrast to the diverse transcriptional responses of *N. stella* to deoxygenation and chemical amendments, *B. argentea* exhibited up-regulation of genes only under the NO_3_^-^ amendment in hypoxia. Specifically, genes related to the assimilatory sulfite reductase (*aSR*), glutathione reductase (*GTR*), and Complex I (*CI*) were up-regulated in the NO_3_^-^ amendment compared to the hypoxia control and the H_2_O_2_ amendment (SI Figure S5). Several carbon metabolism genes, including the mitochondrial *mMDH*, cytosolic *PC*, and peroxisomal *pPGK*, were also upregulated in the NO_3_^-^ amendment compared to the H_2_O_2_ amendment, but not in comparison to the hypoxia control (SI Figure S5). Unlike the active regulation of *Pex* genes in *N. stella*, no differential expression of the *Pex* genes was detected in *B. argentea* under the examined conditions.

## Discussion

Microbial eukaryotes have evolved variable combinations of carbon and energy metabolism in their organelles consistent with their ecological specialization, such as modifications observed in mitochondria (14, 41) and peroxisomes (42, 43). Our (meta)transcriptomic analysis of two cytologically and ecophysiologically distinct foraminifera from the SBB, *N. stella* and *B. argentea* (Table 1), revealed contrasting utilization of these organelles. As expected from the peroxisome-ER complexes observed in previous cytological analyses (7), our analysis showed *N. stella* actively expressed a wide range of genes regulating the function and proliferation of peroxisomes (Figure 1). However, *B. argentea* lacked the expression of some key genes related to peroxisome protein localization (*Pex19*), recycling and degradation of peroxisomal import factors (*Pex2*), and metabolite transport and peroxisome proliferation (*Pex11*), indicating reduced peroxisomal activities consistent with the sparse distribution of peroxisomes in *B. argentea* (Table 1).

The identification of transcripts encoding beta-oxidation and the glyoxylate cycle revealed that *N. stella* may use intracellular lipid droplets as a primary carbon source. This is consistent with the dearth of observed food vacuoles and the abundance of lipid droplets in *N. stella* (23) and in closely related species, such as *Nonionellina labradorica* (25). The predicted integration of beta-oxidation and the glyoxylate cycle in *N. stella* peroxisomes is similar to the glyoxysomes in vascular plants, which are specialized peroxisomes that break down fatty acids to support central carbon metabolism and are linked to the metabolism of mitochondria in germinating seeds (44, 45). In contrast, *B. argentea* lacked the expression of key genes in beta-oxidation and the glyoxylate cycle (Figure 2 **&** 3).

*B. argentea* instead were predicted to employ gluconeogenesis and the oxidative pentose phosphate pathway in the peroxisome (Figure 2 **&** 3), indicating an active central carbon metabolism likely driven by digestion of phagocytosed organic materials. The predicted peroxisomal localization of gluconeogenesis in *B. argentea* could allow the simultaneous carbon metabolism in both the glycolytic and the gluconeogenic directions. This may serve as an alternative to allosteric regulation that prevents toxic accumulation of metabolic intermediates (46), similar to the glycosomes or gluconeosomes described in diplonemids and kinetoplastids (14, 21, 47–49).

Based on predictions of protein localizations, the categorization of *N. stella* and *B. argentea* peroxisomes into a glyoxysome-like and a glycosome-like metabolism, respectively, is consistent with their different cytological traits and ecological distributions (Table 1). The predicted localization of the glyoxylate cycle to the peroxisomes of *N. stella* may represent an adaptation of protistan peroxisomes to use H_2_O_2_ for lipid metabolism in the SBB chemocline. The predicted compartmentalization of glycolysis and gluconeogenesis in *B. argentea* suggests this adaptation could be widespread in microeukaryotes and may reflect convergent evolution associated with hypoxia tolerance (21, 50).

The electron transport system of *B. argentea* and *N. stella* appeared to be an integration of oxygen-dependent and nitrate-dependent metabolisms, likely reflecting adaptations to hypoxic and/or anoxic environments. Diverse electron transport machinery in the mitochondria, including denitrification (23), alternative oxidase (AOX), and sulfide:quinone oxidoreductase (SQOR), was identified among the gene transcripts in either or both foraminifera (Figure 2 **&** SI Figure S1). Genes encoding SQOR have been reported before in other foraminifers, such as *Reticulomyxa filosa, Ammonia* sp., and *Elphidium margaritaceum* (*51*).. The presence of enzymes associated with denitrification has been discussed in depth for both *N. stella* and *B. argentea* (23), as well as in other foraminifers (24, 26). Additionally, the AOX contributes to redox balance by serving as an alternative path to the proton-pumping machinery (Complex III/IV) of the ETC (40), alleviating potential ROS or reactive nitrogen species (RNS) stress caused by the intracellular production of H_2_O_2_, oxygen radicals, and/or nitric oxide from electron leakage of the ETC (52).

Despite oxygen being a common terminal electron acceptor for AOX, genes encoding this protein were expressed across all treatments, including both the anoxic and euxinic conditions (26). Constitutive expression of this gene could have resulted from adaptations to rapid fluctuations of oxygen (53), endogenously sourced oxygen hypothetically produced by catalases in the anoxic treatment (7) or oxygenic denitrification by Nor/Nod (23). Interestingly, recent evidence has shown that nitrite can serve as an alternative electron acceptor for AOX under hypoxia and anoxia in some plants (54), suggesting a potential mechanism for the activation of foraminiferan AOX under anoxia that deserves dedicated biochemical investigation. The observation of SQOR transcripts solely in *N. stella* indicates its capacity to use H_2_S as an electron donor to fuel respiration. The SQOR contributes to a reduced quinol pool that can be used by Nor/Nod to drive denitrification. This is corroborated by the upregulation of denitrification genes in *N. stella* with NO_3_^-^ amendment under euxinia compared to anoxia (SI Figure S4). The expansion of *N. stella*’s electron transport system to utilize H_2_S as an alternative electron donor reflects an adaptive feature for the conservation of carbon sources in *N. stella*, which may be carbon limited as this SBB population of *N. stella* lacks evidence of digestion (23) and utilizes a truncated TCA cycle (Figure 2).

Quinol:fumarate oxidoreductase (QFR) is commonly considered an adaptive mechanism for anaerobic respiration in eukaryotic organisms under anoxic conditions (38). We found constitutive expression of genes encoding QFR solely in *N. stella* (Figure 4B), suggesting potential adaptations of this species to an anaerobic lifestyle. The conservation of QFR based on the presence of a glycine residue in the large cytochrome subunit of foraminifers (Figure 4A) indicates an ancestral state of this taxon that may have evolved in an anoxic milieu. In *N. stella*, transcripts encoding RQUA of the rhodoquinone biosynthesis pathway further supports the potential presence of fumarate reduction (51, 55). Interestingly, transcripts of succinate dehydrogenase (SDH) were observed in both *N. stella* and *B. argentea.* The SDH formed a monophyletic group with the QFR, suggesting that both the QFR and SDH may have originated from a common foraminiferal ancestor capable of fumarate reduction. Given the prevalence of QFR in other foraminifera and the expression of the RQUA gene in *B. argentea*, a QFR could be encoded in the *B. argentea* genome, although it was not detected in the (meta)transcriptomes represented in this study.

*B. argentea* additionally demonstrated potential capacities for substrate-level phosphorylation through the coupling of succinate:acetate CoA transferase (SCoAT) and succinyl-CoA synthetase (SCS). Genes encoding SCS and SCoAT have been previously identified in related species (26), but this is the first documentation of the expression of the *SCoAT* gene in foraminifera. Carbon conservation was inferred from the predicted mitochondrial PFOR, the cytosolic PNO, and the cytosolic acetyl-CoA ligase (ACoAL), another gene previously not documented in foraminifera (Figure 2). The utilization of substrate-level phosphorylation and the constitutive expression of denitrification pathways by *B. argentea* under hypoxia indicated the potential lifestyle of a facultative anaerobe. The up-regulation of *aSR* and *GTR* by *B. argentea* in hypoxia NO_3_^-^ amendment (SI Figure S5) was consistent with an increased tolerance of ROS stress for maintaining active carbon metabolism (56, 57).

In contrast, *N. stella* demonstrated a wider range of metabolic responses to the H_2_O_2_, NO_3_^-^, and H_2_S amendments under both hypoxia and anoxia by using a combination of peroxisomal and mitochondrial pathways. The H_2_O_2_ amendment induced the up-regulation of genes associated with beta-oxidation and the glyoxylate cycle in *N. stella*, likely linked to the utilization of intracellular lipid droplets that are abundant in this species (23). The release of carbon substrates by beta-oxidation and the glyoxylate cycle appears to be coupled with carbon storage via gluconeogenesis (i.e. up-regulation of *FBP*) and a more active electron transport system, based on the up-regulation of the denitrification and *SO* genes (Figure 5A). The H_2_O_2_ amendment also stimulated up-regulation of ATP-producing Complex V genes, which was consistent with the experimentally measured increases in intracellular ATP production of *N. stella* with the addition of H_2_O_2_ (7). The metabolism of *N. stella* under the H_2_O_2_ amendment may reflect its native metabolic state in the SBB sediments, as no differential gene expression was identified between the H_2_O_2_ amendments under anoxia or hypoxia versus their corresponding control (without amendment) conditions (Figure 5A **& SI Dataset S2**).

The NO_3_^-^ amendment of *N. stella* induced differential gene expression compared to both the control and the H_2_O_2_ amendment. The down-regulation of *FBP* and the up-regulation of *FBA* in anoxia NO_3_^-^ amendment may indicate the activation of carbon metabolism in the glycolytic direction, which can be an adaptive mechanism in response to the lack of carbon substrates due to down-regulation of the beta-oxidation (Figure 5B). The up-regulation of both *RQUA* and Complex II genes indicates the potential activation of fumarate reduction. This is coordinated with the down-regulation of denitrification genes and likely reflects a metabolism limited by electron donors, as denitrification competes with fumarate reduction for the quinone pool. Further evidence for limited availability of electron donors was observed under the anoxic NO_3_^-^ amendment in comparison to the euxinic NO_3_^-^ amendment. With H_2_S as an alternative electron donor via SQOR, *N. stella* showed an up-regulation of genes involved in denitrification, beta-oxidation, the glyoxylate cycle, and gluconeogenesis (SI Figure S4). This reflects an adaptive feature for the conservation of carbon sources in *N. stella*, which may be carbon limited as this species lacks evidence for digestion through the presence of food vacuoles (23). Lastly, the up-regulation of ROS response genes (e.g. *APX*, *GTP*) in the NO_3_^-^ amendment of *N. stella* likely indicates cellular mitigation of the effects of electron leakage during fumarate reduction (58).

The evolution of mitochondria and peroxisomes are often coupled in eukaryotes (17, 18, 22) and represents a spectrum of metabolic phenotypes (13, 14). Combinations of metabolic pathways known in MROs were predicted to localize in the mitochondria of *N. stella* and *B. argentea* (Figure 2); however, mitochondria of these foraminifer still maintain the oxygen-dependent ETC (CIII, CIV, AOX) (Figure 2) and may perform anaerobic respiration through denitrification (23). These unique metabolic capabilities have not been identified in any other protists and may allow our two benthic foraminifera to survive in both oxygen depleted and aerated conditions. The organelle-specific metabolic flexibility documented in this study offers a contrast to the mitochondrial metabolism of conventional MROs found in anaerobic protists. These differences suggest that the mitochondria of these two benthic foraminifera have evolved specialized metabolic tools that support their ability to respond to varied oxygen concentrations and milieu chemistry. The plasticity observed in the mitochondria was enhanced by the peroxisomes in both species, which are metabolically distinct not only from each other, but also from known anaerobic peroxisomes (17, 18, 22). The selective pressures associated with survival in highly variable habitats have conferred our two foraminifera with diverse metabolic toolkits of mitochondrial and peroxisomal metabolism that enables survival in SBB chemoclines.

Overall, the two foraminifers demonstrated distinct metabolic adaptations to hypoxia and anoxia, and had different capacities to use compounds often abundant in SBB pore water (4, 23). This indicates benthic foraminiferal metabolism is not uniform across species. While the ability to perform denitrification and anaerobic energy metabolism are established for *N. stella* (4, 23), we show here that *N. stella* of the SBB are adept at surviving anoxia, having metabolically flexible mitochondria that function in coordination with the predicted metabolism of its peroxisomes. The metabolism predicted for *B. argentea* suggested the capacity to digest food vacuole contents as a facultative anaerobe. These species represent only two examples of the unique cytologies that occur along the SBB chemocline. Additional cytological diversity is known among other foraminifera, such as *Bulminella tenuata*, an endobiont-bearing species that also exhibits close associations between mitochondria and peroxisomes (7), or an as-yet undescribed agglutinated foraminifer with coccoid endosymbionts that lacks peroxisome-ER complexes in favor of few peroxisomes (28, 59). The highly variable cytology observed among foraminifera of the SBB chemocline and elsewhere, such as hydrocarbon seeps (60), implies potential distinctions in their metabolic capacity, which may influence biogeochemical cycling and calcite biomineralization, as the presence of symbionts likely shifts **δ**^13^C_calcite_ values in one foraminifera species (61). Further understanding the highly variable metabolic adaptations that can occur across foraminifera will allow for more informed reconstructions of paleoceanographic conditions and strengthen biogeochemical and ecological predictions of shifts in organismal and metabolic niches under future ocean deoxygenation. Lastly, these findings provide additional insights into the biogeochemical roles and the evolution of this ecologically significant protistan taxon that diversified in the Neoproterozoic.

## Materials and Methods

### (Meta)Transcriptome data

Transcriptome and metatranscriptome data were obtained from the NCBI BioProject PRJNA714124. Details of the experimental settings were described in reference (23). Briefly, surface sediments containing either *N. stella* or *B. argentea* of the Santa Barbara Basin were subjected to laboratory treatments under four oxygen conditions (i.e. aerated, hypoxic, anoxic, and euxinic) and four chemical amendments: the no amendment (Control), nitrate (NO_3_^-^), hydrogen peroxide (H_2_O_2_), nitrate and hydrogen peroxide (NO_3_^-^ + H_2_O_2_), with 2-3 replicates in ∼75 cm^2^ polypropylene containers for each species given each combination of the oxygen and amendment conditions. Following a treatment period of three days, *N. stella* or *B. argentea* were picked into pools of 25 (cleaned) conspecifics, and each pool was counted as a biological replicate for transcriptomic or metatranscriptomic sequencing using the PolyA- or Total RNA-based library preparation, respectively (23). Bar-coded library samples were sequenced using two sequencing runs on an Illumina NovaSeq platform for the polyA selected mRNA library and two sequencing runs on an Illumina NextSeq platform for the total RNA library. Overall, between 3-7 biological replicates per amendment/treatment were analyzed, with each biological replicate represented by pooling all sequences from its two sequencing runs. Detailed information of all samples referenced in this study and their corresponding data in the National Center for Biotechnology Information (NCBI) Sequence Read Archive (SRA) database are summarized in **SI Dataset S3**.

### Transcriptome assembly and annotation

The *N. stella* and *B. argentea* transcriptomes and metatranscriptomes were quality checked and processed as described previously (12). Briefly, reads were quality checked with TrimGalore (62), a wrapper script built around Cutadapt (63) and FastQC (64). Each (meta)transcriptome was assembled individually using the *de novo* transcriptome assembly functions in rnaSPAdes v3.14.1 with default parameters (65).

Due to the large size of the nucleotide dataset (greater than fifty million contigs), the lin-clust clustering algorithm from MMSEQS2 was used to cluster nucleotide sequences with a minimum sequence identity of 97% using default settings (66). Coding sequences (CDS) within a representative contig from each cluster were identified using geneMarkS-T with default settings (67) and the protein sequences encoded by the resulting CDS were further clustered using the Cluster Database at High Identity with Tolerance (CD-HIT) (68) with a minimum sequence identity of 99% and 95% to identify representative sequences for further annotations and taxonomic assignments, respectively. Representative protein sequences from the 99% CD-HIT clustering were annotated with EggNOG V.2 (69) to assign enzyme commission (EC) numbers. The EggNOG annotation did not capture all subunits of mitochondrial electron transport chain complexes. Therefore, targeted identification of these subunits was performed by examining the presence and organization of protein domains based on matches to hidden markov models (HMM) from the Pfam 27.0 database (70) using HMMER 3.1 (71) with an independent e-value cutoff of < 1E-5. Genes identified as missing or as expressed in only one species were further searched for using a homology-based approach by mapping reference proteins against the assembled contigs using tblastn (72). Three sets of protein sequences were used for this search: foraminifera proteins identified from the automated prediction of coding sequences for genes that were missing in one species, sequences of the ETC complexes from the Protein Data Bank (PDB IDs: SRFR, 3VRB, 1NTZ, and 5B3S for CII, CII, CIII, and CIV respectively), and the RQUA sequence from *Reticulomyxa filosa* (Accession: ETO28810.1). Sequences that shared 60% identity and 70% coverage to the reference proteins and were not already identified in the CDS predictions were added to the list of annotated genes. Two RQUA transcripts found through this search were included in the analysis despite sharing a sequence identity of 47.6% and 45.5% to the RQUA sequence from *Reticulomyxa filosa* based on the identification of binding sites known for RQUA (Figure 4C).

Representative protein sequences from the 95% CD-HIT clustering were mapped against the NCBI non-redundant protein database using the DIAMOND BLASTP (73). If a sequence had an alignment to foraminifera out of the top 25 best hits, it was taxonomically assigned to the foraminifera group. If a representative protein sequence had a majority of the top 25 hits assigned to diatoms, other eukaryotes, bacteria, or archaea, the sequence was taxonomically assigned based on the majority mapping.

### Organelle localization predictions

Peroxisome localization was predicted using a consensus approach that combined motif-based identification of targeting signals with homology-based search of well-curated peroxisomal proteins (SI Figure S6). The *Pex5*, *Pex7*, and *Pex19* genes were identified in the foraminiferal (meta)transcriptomes and the Pex5, Pex7, and Pex19 proteins recognize peroxisome targeting signal 1 (PTS1), peroxisome targeting signal 2 (PTS2), and peroxisomal membrane protein import signal (PMP), respectively. Correspondingly, these three signals were used to predict peroxisomal localization (**SI Table S1**). The predictions were informed by a consensus of four strategies (SI Figure S6), using tools from the MEME Suite, version 5.1.1 (74). Each protein sequence identified from the (meta)transcriptome was analyzed using the motif-based approach, where a consensus score from one to four was assigned for each of the peroxisomal import signals by counting the number of strategies that resulted in a motif match to the target sequence. Confidence levels of peroxisome localization prediction were evaluated based on the number of strategies that support the motif-based mapping and a homology search to the peroxisomeDB reference database, a database of peroxisomal proteins from diverse organisms, including protists (75), using DIAMOND BLASTP (73) (**SI Table S2**).

Mitochondrion localization was predicted based on homology and a consensus of three predictors: MitoFates version 1.2, TargetP version 2.0, and PredSL (76–78), combinations of which have been utilized with success in putatively anaerobic protists previously (79, 80). Combined probabilities of mitochondrion localization were calculated by multiplying the probabilities from all three predictors to provide an aggregated estimation. In parallel, DIAMOND BLASTP (73) was used to identify homology from our assembly to a reference database consisting of mitochondrial proteins from OrganelleDB (81), Mitocarta (82), the *Acanthomoeba castellanii* mitochondrial proteome (83), the *Andalucia godoyi* proteome (84), and mitochondrial sequences from the plant proteome database (85). Confidence levels were assigned to the putative mitochondrial proteins by integrating the combined probability of localization and the homology identifications to the reference mitochondrial protein database (**SI Table S2**).

Taxonomic assignments were additionally used for the filtering of peroxisomal and mitochondrial pathways. Only proteins taxonomically assigned to foraminifera or other eukaryotes were included in the metabolic reconstruction. A number of metabolic processes were placed to mitochondria because they represent known or proposed genes in mitochondria. These included subunits of the electron transport chain complexes and mitochondrial transporters (**SI Dataset S1**) identified through homology to the above-mentioned mitochondrial protein reference database and the transporter classification database (TCDB) (86) using DIAMOND BLASTP with a sequence identity cutoff of at least 30% and a reference coverage cutoff of at least 50%.

Similarly, several proteins were assigned to the peroxisome despite the absence of signal motifs, including Pex proteins, transmembrane transporters, catalase and peroxidase, and Isocitrate lyase (**SI Dataset S1**). Isocitrate lyase was included due to homology to known peroxisomal isocitrate lyases. Previous research also showed that isocitrate lyase proteins without import signals may be able to localize to the peroxisome by co-importing with other proteins (87). Catalases are known to have a modified peroxisomal import signal, which may explain why they were not identified in our analysis (88). The catalase and peroxidase proteins are commonly known to localize in the peroxisome. Further, the peroxisomal localization of catalase in *N. stella* is seen through the presence of the crystalline cores formed by the catalase aggregates in its peroxisomes (7). Transport of small soluble metabolites such as malate and succinate across the peroxisomal membrane was included in our prediction, as the transport of small metabolites is commonly mediated by a variety of non-selective channels through which these compounds can freely diffuse (89). Specific transporters for cofactors such as NAD, CoA, ATP, acyl-CoA, and fatty acids were identified through homology searches to known peroxisomal transporters from the Swiss-Prot database, release-2021_01 (90), and the peroxisomeDB using DIAMOND BLASTP (73) with a sequence identity cutoff of at least 30% and a reference coverage cutoff of at least 50%. While little support was found for the localization of the denitrification proteins (pNR, NirK, and Nor/Nod) to the mitochondria, its involvement in respiration may require an association with the mitochondria. This was proposed in previous studies by our group and others (23, 24), so these proteins were included in our reconstruction of mitochondrial metabolism.

Proteins not localized to peroxisome or mitochondrion were classified as cytosolic and were included in the metabolic reconstruction for the completion of central carbon metabolism. Metabolic connections among the peroxisomal, mitochondrial, and cytosolic compartments were examined by tracing the carbon flow in the metabolic network using the *findprimarypairs* function in the PSAMM software package version 1.0 (91).

### Identification of succinate dehydrogenase/quinol:fumarate oxidoreductase

Binding sites relevant to the function of Complex II were analyzed based on multiple sequence alignments (MSA), generated using the PROfile Multiple Alignment with predicted Local Structures and 3D constraints (PROMALS3D) (92). Each subunit of Complex II was aligned to reference sequences from the protein data bank, marine microbial eukaryotic transcriptome sequencing project (MMETSP) (93), Swiss-Prot (90), NCBI RefSeq non-redundant protein database (94), and from eukaryotes known to live in anoxic conditions (58). Three-dimensional protein structures of the Complex II from *Ascaris suum* (PDB ID 3VR8), *Sus scrofa* (PDB ID 3FSD), and *Gallus gallus* (PDB ID 2H89) were used in PROMALS3D (92) to guide the MSA construction for each subunit. Alignments of the four core subunits (CII-SDHA, CII-SDHB, CII-SDHC/QFRC, CII-SDHD) were concatenated and used as input for the phylogenetic reconstruction using RAxML version 8.2 (95). A concatenated alignment of all subunits resulted in an alignment with 39.28% gaps and 1677 distinct alignment positions. No trimming was performed on the MSA. An optimal substitution model was selected by RAxML using the automatic model selection under Akaike Information Criterion (AIC) (96). The best model, LG substitution with the GAMMA model of rate heterogeneity and empirical base frequencies, was applied for the phylogenetic reconstruction of Complex II using 100 bootstraps. Specific amino acid residues relevant to the functions of QFR and SDH were identified following previous studies on the quinone-binding pocket of these proteins (38).

### Differential expression analysis

Differentially expressed genes were identified using representative transcripts from the assembly after clustering at 99% minimum amino acid sequence identity (described above). Quality controlled sequencing reads from each sample were mapped to these representative transcripts using BBMap from the BBTools package (97). Briefly, read mapping files from each sample were sorted, indexed with the SAMtools package version 1.7 (98), and used for the calculation of transcript abundance using the Pileup function in BBTools version 37.36 (97). The species-specific mapping of transcripts was achieved by counting the abundance of a transcript across all treatments of a species. A denoising step was applied to remove sporadic mappings of reads by exclusively assigning each transcript to a species (i.e. *N. stella* or *B. argentea*) if the read mapping was dominated (e.g. with a 10 times higher abundance) by samples from that species. If a transcript received a similar number of mapped reads from both species, the transcript was assigned to both species if > 60 reads were mapped across all samples of each species. Transcripts mapped to < 60 reads in both species were removed from further consideration, as having < 60 reads would on average represent only around one read mapped per sample in each species. Count data were summed for all transcripts mapped to the same species and annotated as the same gene, and this combined count data were used as inputs for the differential gene expression analysis using DESeq2 version 1.26.0 in R (99). Each combination of chemical amendments and oxygen conditions was used as an experimental treatment for the modeling of differential gene expression. Additionally, a variable was introduced to the statistical model with two factor levels to control for the potential variation introduced from the use of two different library preparation techniques: (1) polyA selection that targets the polyadenylated mRNA from eukaryotes, and (2) total RNA preparation that targets all mRNA molecules from eukaryotes and prokaryotes (23). Statistical significance was based on the Wald test by estimating variation in log-fold change (LFC) using differences between experimental treatments. Results from the Wald test were further considered if the two sets of samples represented the same oxygenation treatment but different chemical amendment, or if they represented the same chemical amendment but different oxygenation treatments. For example, the non-amended hypoxic treatment and the NO_3_^-^ amended hypoxic treatment reflected a comparison of chemical-driven responses. In contrast, the NO_3_^-^ amended hypoxic treatment and the NO_3_^-^ amended anoxic treatment reflected a comparison of oxygen-driven responses. A significance threshold <0.05 was used on the adjusted p-values based on a Benjamin Hochberg correction for false discovery rates. The expression of genes in a treatment was calculated by taking the mean of the median ratio normalization (MRN) values across all biological replicates of a given treatment followed by log_2_ transformations. Heatmaps were created based on the log_2_ transformed MRN values using ggplot2, version 3.3.2 (100). Differential expression of genes was similarly plotted using ggplot2.

## Data Availability

Raw sequencing data referenced in this study was obtained from the NCBI database under BioProject accession PRJNA714124. Detailed reference of the Sample, Experiment, and Run numbers are listed in **SI Dataset S3** of this manuscript. Data files generated from the computational workflow will be released via a public FigShare data repository. A private link for access by reviewers can be accessed at https://figshare.com/s/10e011de3338eef270bc. All software and algorithms applied in this study are documented in the Methods section. A public git repository documenting the computational workflow will be available at https://github.com/zhanglab/SBB_Foram_MP_Analysis/.

## Author Contributions

J.M.B., Y.Z., C.M.H. and V.P.E. conceived the study. J.M.B., D.J.B., C.M.H., S.D.W., and H.L.F. contributed to sample collection. F.G., J.M.B., and D.J.B. performed the experiment. C.P., Y.Z., F.G., E.B.B., D.R.U., S.D.W, C.M.H., and J.M.B. analyzed and interpreted the data. C.P., Y.Z., F.G., and J.M.B. composed the manuscript. All authors gave feedback on the manuscript.

## Funding

This project was funded by the U.S. National Science Foundation IOS 1557430 and 1557566. H.L.F. acknowledges support from the Swedish Research Council VR (grant number 2017-04190). C.P. acknowledges partial support from the U.S. National Science Foundation DBI 1553211. E.B.B. acknowledges support from the URI MARC U*STAR grant from the National Institute of General Medical Sciences of the National Institutes of Health under grant number T34GM131948 (Niall G. Howlett, PI)

## Competing Interests

The Authors declare no competing interests.

## Supporting Information (SI)

### SI Figures

**SI Figure S1:**
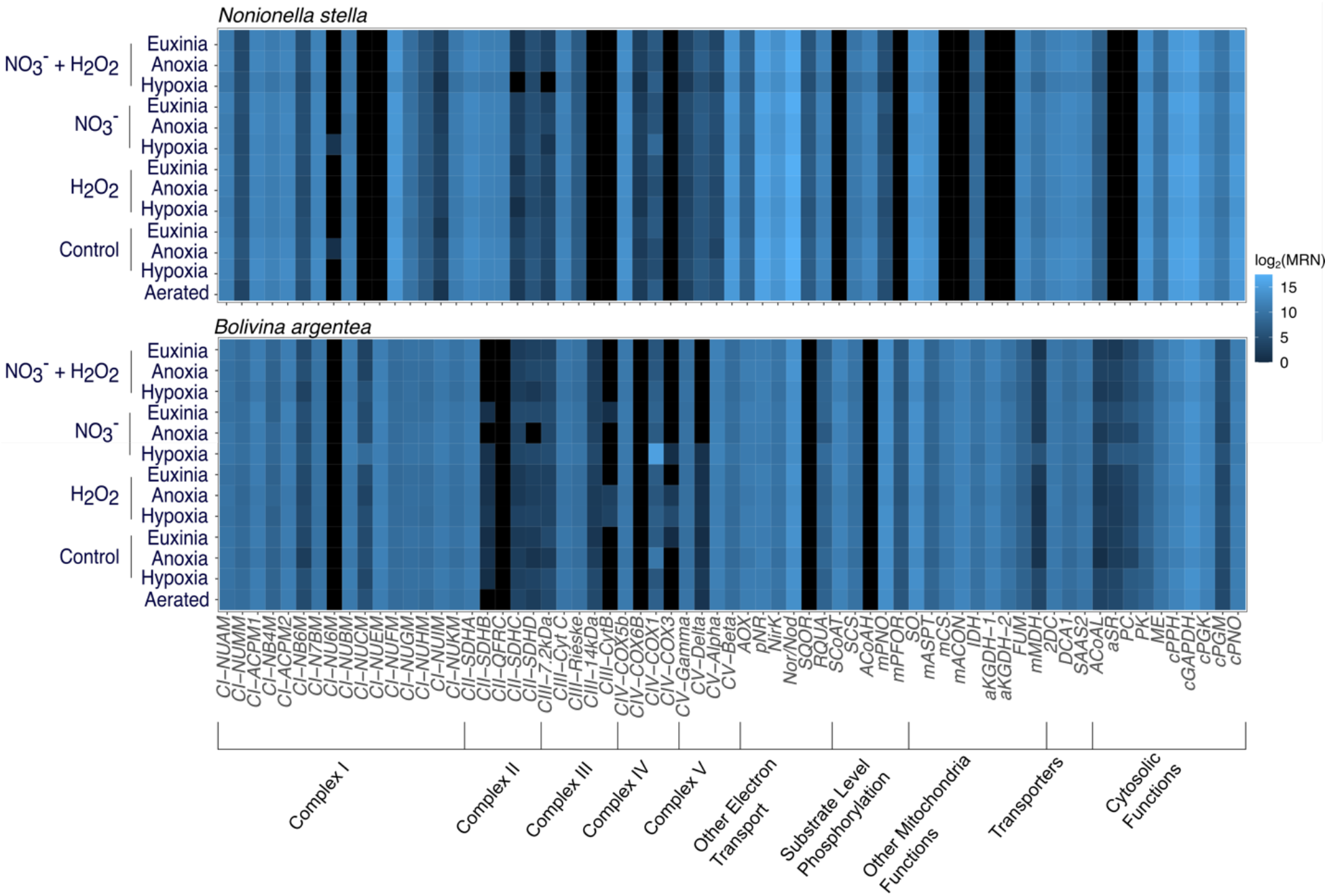
Heatmaps by species showing the log_2_ transformed median ratio normalization (MRN) values for all genes that encode mitochondrial and cytosolic proteins. Black indicates no expression. Abbreviations align with those in **SI Table S1.**

**SI Figure S2:**
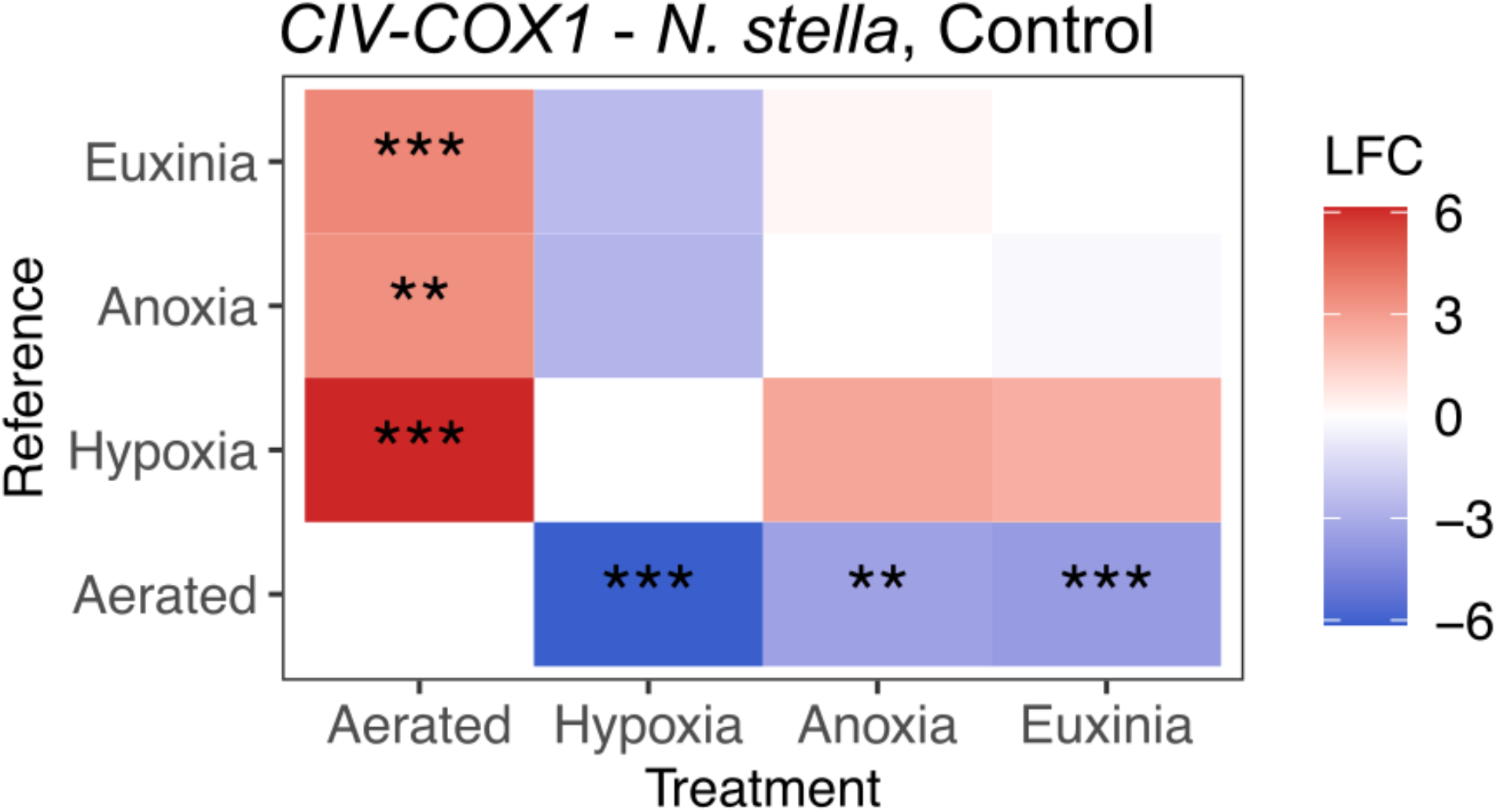
Matrix of differential expression of genes encoding the *COX1* subunit of Complex IV across treatments with no chemical amendment (Control). The pairwise comparisons are color coded based on log_2_ fold change (LFC) values. * indicates adjusted p-value < 0.05. ** indicates adjusted p-value < 0.01. *** indicates adjusted p-value < 0.001.

**SI Figure S3:**
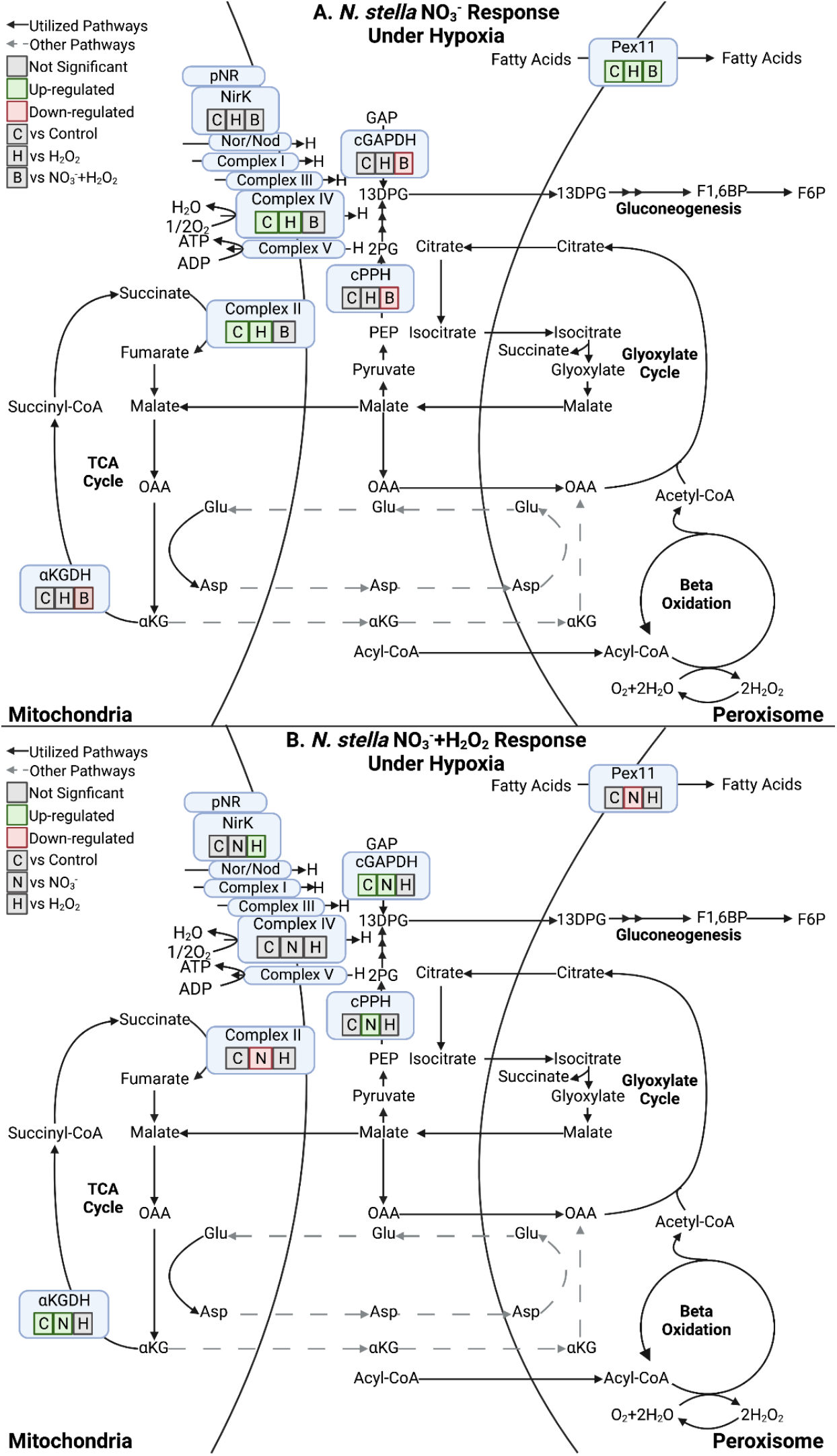
Metabolic responses of *N. stella* to different chemical amendments under hypoxia. **A.** Differential expression of genes under the NO_3_^-^ amendment in contrast to the control (C), H_2_O_2_ (H), and combined NO_3_^-^-+ H_2_O_2_ (B) amendments. **B.** Differential expression of genes under the combined NO_3_^-^-+ H_2_O_2_ amendment in contrast to the control (C), NO_3_^-^- (N), and H_2_O_2_ (H) amendments. Significant differential gene expressions between contrasts (adjusted p-value < 0.05) are highlighted in green (Up-regulated) or red (Down-regulated). Contrasting conditions with no significant differential gene expressions are indicated in gray (Not Significant).

**SI Figure S4:**
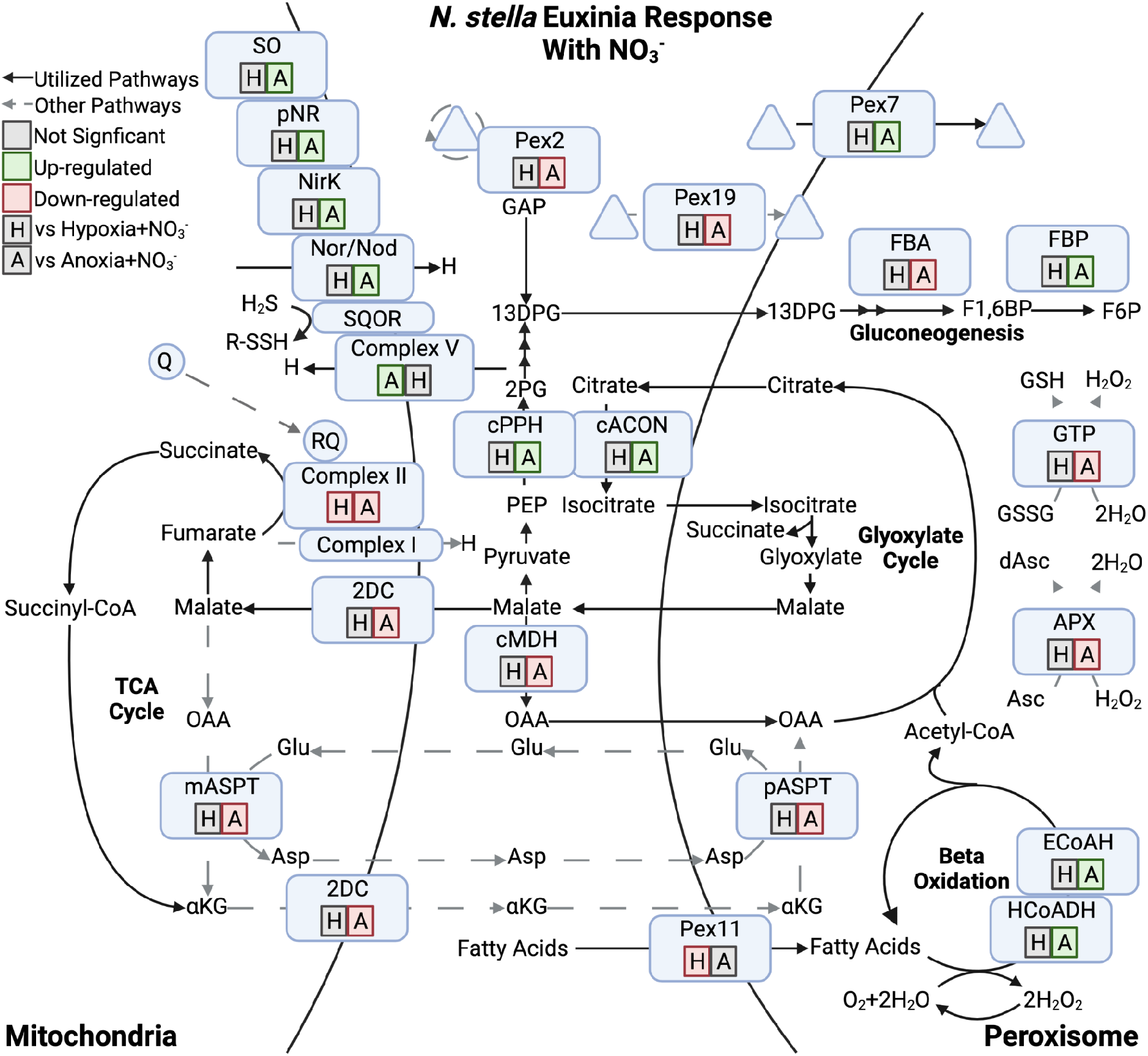
Metabolic response of *N. stella* to the euxinia condition in contrast to hypoxia (H) and anoxia (A) under the NO_3_^-^ amendment. Triangles represent transported and recycled proteins. Significant differential gene expressions between contrasts (adjusted p-value < 0.05) are highlighted in green (Up-regulated) or red (Down-regulated). Contrasting conditions with no significant differential gene expressions are indicated in gray (Not Significant).

**SI Figure S5:**
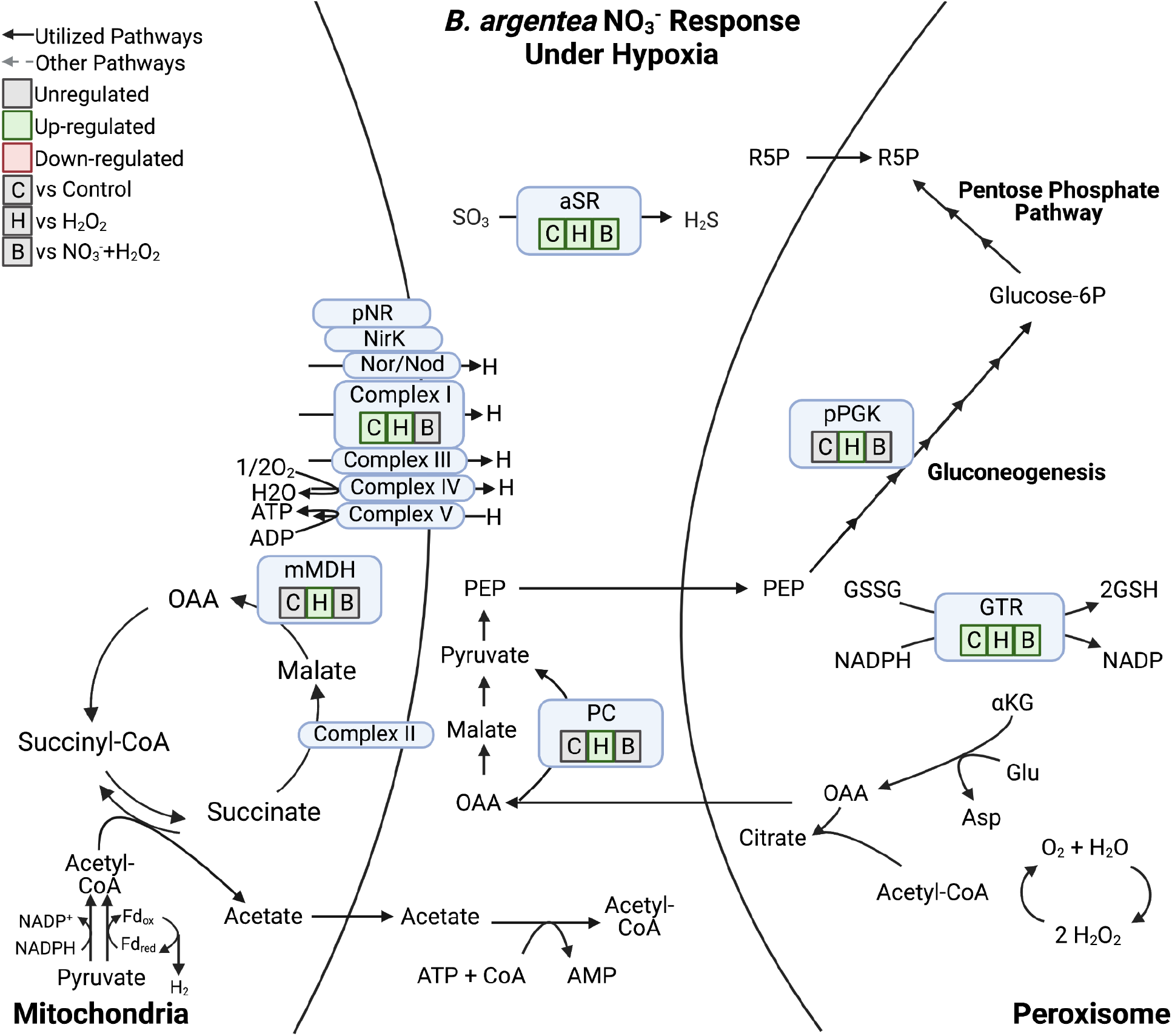
Metabolic response of *B. argentea* to the NO_3_^-^ amendment in contrast to the control (C), H_2_O_2_ (H), and combined NO_3_^-^+ H_2_O_2_ (B) amendments under hypoxia. Significant differential gene expressions between contrasts (adjusted p-value < 0.05) are highlighted in green (Up-regulated) or red (Down-regulated). Contrasting conditions with no significant differential gene expressions are indicated in gray (Not Significant).

**SI Figure S6:**
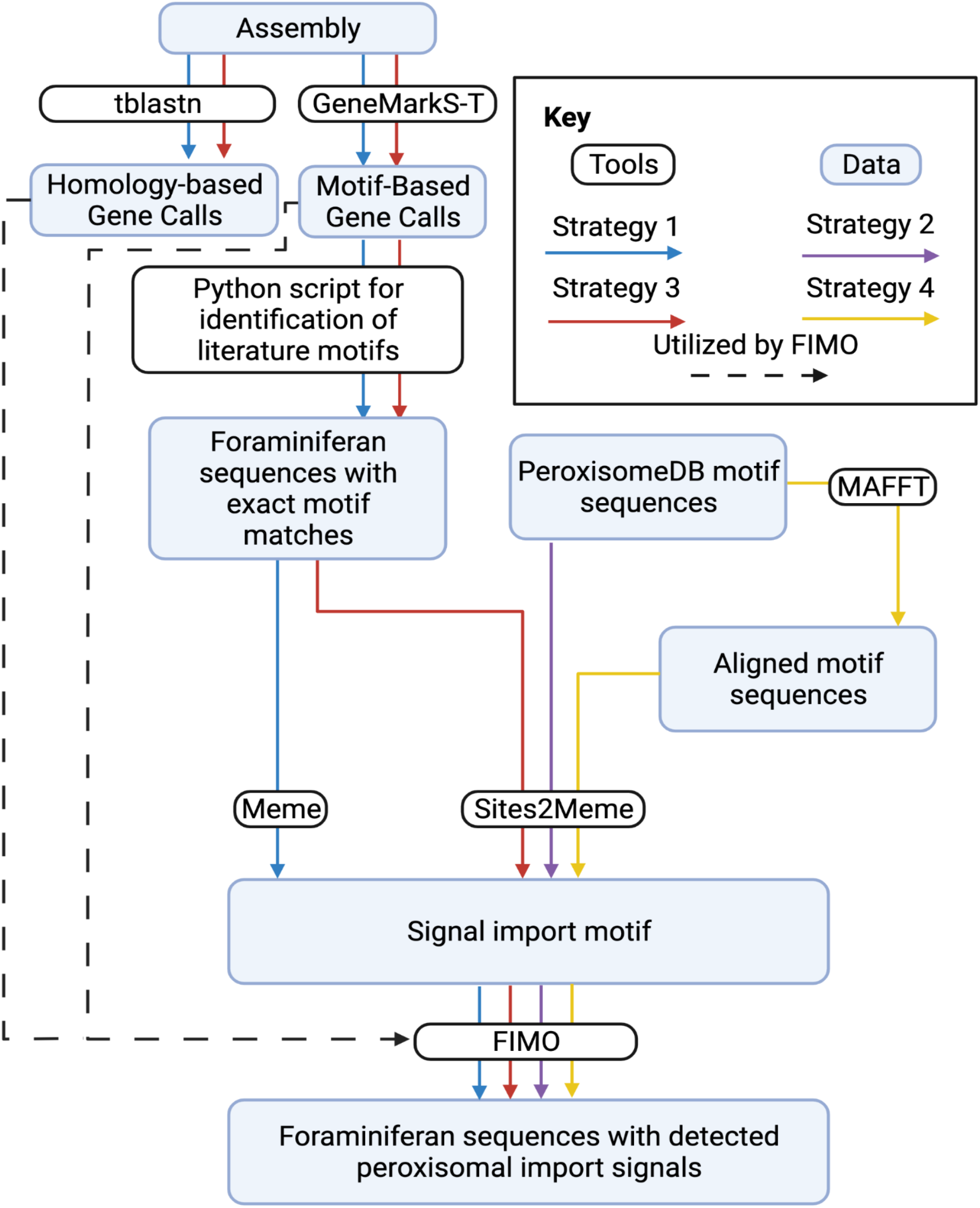
Flowchart showing the consensus of four strategies implemented in this study for the identification of peroxisomal-localization motifs based on protein sequences annotated from the foraminiferan transcriptome and metatranscriptome assembly.

### SI Tables

**SI Table S1:** List of genes identified in this study. Compartment indicates predicted localization for each gene. Gene association indicates identifiers of contigs that encode a corresponding gene within each species. Confidence of localization indicates the confidence of localization to the mitochondrion or the peroxisome (**SI Table S5)** based on a consensus approach specified in the **Materials and Methods**. Log_2_(MRN) indicates the log_2_ transformed median ratio normalization (MRN) values used for the visualization of gene expression levels in heatmaps (**Materials and Methods**).

**SI Table S2:** List of differentially expressed genes identified based on the DESeq2 differential gene expression analysis. Treatment and reference columns indicate the comparison being made for each gene. Basemean represents the mean expression of the gene across all treatments. Log_2_ fold change shows the log_2_ fold change of expression in the treatment as compared to the reference. Log_2_ fold change SE is the standard error in the log_2_ fold change. Stat indicates the Wald statistic. P-value indicates the significance before corrections for false discovery rates. Adjusted p indicates the p-value after applying a Benjamin Hochberg correction for false discovery rates.

**SI Table S3:** National Center for Biotechnology Information (NCBI) Sequence Read Archive (SRA) accession numbers, species, oxygen treatment, chemical amendment, and type of library prep organized by sample ID. Each row represents a (meta)transcriptome determined from a biological replicate of 25 pooled cells.

**SI Table S4:** Reference motifs used for the identification of genes localized to the peroxisome: peroxisome targeting signal 1 (PTS1), peroxisome targeting signal 2 (PTS2), and peroxisomal membrane proteins (PMP).

**SI Table S5:** Confidence levels defined in the consensus-based prediction of peroxisomal or mitochondrial protein localizations.

